# Effects of experiencing CS-US pairings on instructed fear reversal

**DOI:** 10.1101/2022.04.05.487162

**Authors:** David Wisniewski, Senne Braem, Carlos González-García, Jan De Houwer, Marcel Brass

## Abstract

Fear learning allows us to identify and anticipate aversive events, and adapt our behavior accordingly. This is often thought to rely on associative learning mechanisms where an initially neutral conditioned stimulus (CS) is repeatedly paired with an aversive unconditioned stimulus (US), eventually leading to the CS also being perceived as aversive and threatening. Importantly, however, humans also show verbal fear learning. Namely, they have the ability to change their responses to stimuli rapidly through verbal instructions about CS-US pairings. Past research on the link between experience-based and verbal fear learning indicated that verbal instructions about a reversal of CS-US pairings can fully override the effects of previously experienced CS-US pairings, as measured through fear ratings, skin conductance, and fear-potentiated startle. However, it remains an open question whether such instructions can also annul memory traces in the brain. Here, we used a fear reversal paradigm in conjunction with representational similarity analysis of fMRI data to test whether verbal instructions fully override the effects of experienced CS-US pairings in fear-related brain regions or not. Previous research suggests that only the right amygdala should show lingering representations of previously experienced threat (a so-called “Pavlovian trace”). Unexpectedly, we found evidence for the residual effect of prior CS-US experience to be much more widespread than anticipated, in the amygdala but also cortical regions like the dorsal anterior cingulate or dorso-lateral prefrontal cortex. This finding shines a new light on the interaction of different fear learning mechanisms, at times with unexpected consequences.

## 1. Introduction

Animals developed the adaptive ability to relate stimuli (CSs) with harmful events (USs), which allows them to anticipate and avoid such events in the future (Pavlovian conditioning, Maren, 2001; Öhman and Mineka, 2001). A fundamental advantage of human fear learning is the ability to also learn from verbal instructions (Olsson and Phelps, 2007). In most cases, a single instruction is enough to lead to the desired behavior, without the need for repeated trial and error learning (“Never leave electrical appliances near your bathtub”). Recently, the interaction between associative learning and verbal instructions, and whether verbal instructions can fully override experience-based learning, have received increased attention (Koban et al., 2017; Mertens et al., 2018; Atlas, 2019). Recent studies have shown that behavioral conditioned responses can be fully reversed by merely verbally instructing a reversal of CS-US contingencies (Atlas et al., 2016; Mertens and De Houwer, 2016; Lonsdorf et al., 2017; Atlas and Phelps, 2018). Across these studies, merely instructing the discontinuation of an established CS-US pairing was shown to immediately result in a substantially reduced fear response.

Of all brain regions implicated in processing fear-relevant stimuli (Fullana et al., 2015), the amygdala seems to have a specific role in experienced CS-US pairings (Atlas et al., 2016; Braem et al., 2017). While instructed contingency reversal seems to fully override previously learned responses in most fear-relevant brain regions (Atlas, 2019), the right amygdala shows lingering effects of prior CS-US experience above and beyond the effects of verbal instructions (“Pavlovian trace”, Braem et al., 2017). Although this demonstrates the limits of the effects of verbal instructions on prior CS-US experience, we know much less about the opposite effect of CS-US experience on verbal instruction implementation in fear reversal. One study has shown that instructed fear reversal effects on behavioral and psychophysiological measures can be fully explained through verbal instructions only, with no additional effect attributable to CS-US experience (Mertens and De Houwer, 2016). Our goal was to test whether similar effects can be shown on the neural coding of fear-relevant stimuli as well, which remains unknown.

Previously, neural signals associated with experienced and instructed CS (CS+E) and merely instructed CSs (CS+I) were dissociated to identify Pavlovian traces in the right amygdala (Braem et al., 2017). This study used a static design with constant CS-US associations, and it remains unclear whether such effects generalize to more dynamic reversal learning settings. Another fMRI study implemented a dynamic fear reversal learning paradigm to investigate experience-instruction interactions (Atlas et al., 2016). Here, the authors compared a condition in which participants relied on both experience and verbal instructions, with a condition in which participants relied on experience alone. Keeping experience constant across conditions while manipulating the presence of verbal instructions is a good way to demonstrate effects of verbal instructions on experience-based learning. Yet it is not optimal to demonstrate specific effects of prior CS-US experience on verbal reversal instruction implementation, which requires keeping instructions constant across conditions and manipulating the presence of CS-US experience.

Here, we used a fear reversal paradigm, using representational similarity analysis of fMRI data (Kriegeskorte et al., 2008) to measure neural coding of fear-relevant stimuli (Visser et al., 2013; Braem et al., 2017). Participants first performed a conditioning phase, where they received instructions about safe (CS-, never followed by US) and threatening (CS+, potentially followed by US) stimuli. Crucially, only some CS+s were actually followed by an aversive electrical stimulus (CS+E), while others were not (CS+I). After subsequent verbal reversal instructions, each CS was presented again in a second instructed reversal phase (which we will call ‘reversal phase’ from now on). In line with prior findings, we hypothesized that verbal instructions would be able to fully reverse both the behavioral and neural expressions of previously learned CS-US relations (Mertens and De Houwer, 2016). Only the (right) amygdala, we reasoned, could show effects of prior experience after verbal reversal instructions.

## 2. Methods

### 2.1 Participants

42 participants took part in the experiment (23 female, 19 male, mean age = 23.0, age range = 18 – 34, right-handed, no history of neurological or psychiatric disorders). All participants volunteered to participate and gave written informed consent, had normal or corrected-to-normal vision, and received 40€ for participation. The experiment was approved by the ethics committee of the Ghent University Hospital (registration number B670201421176). Two participants showed excessive head movements inside the MR scanner (>5 mm) and were excluded from all further analyses. The final sample consisted of 40 participants (21 female, 19 male, mean age = 22.9, age range = 18 - 34).

### 2.2 Procedure

#### 2.2.1 Work-up

After entering the MR scanner, an experimenter attached the electrode either to the participant’s right or left leg, and the work-up procedure started. The electrotactile pain stimulus was delivered through a surface electrode (Speciality Developments, Bexley, UK), placed over the ankle (retromalleolar course of the sural nerve). The stimulated leg (left or right) was counterbalanced across participants. Stimulation was administered using a DS5 electrical stimulator (Digitimer, Welwyn Garden City, UK), and stimulation intensity was determined using an adaptive work-up procedure. Individual pain thresholds were determined using an interleaved staircase procedure (Braem et al., 2017), consisting of a total of 20 trials. On each trial, an electrical stimulus was administered, and the participant was asked to verbally indicate pain intensity on a scale from 0 (no pain) to 10 (extreme pain). The 20 trials were randomly divided into two staircase sequences of 10 trials. Each staircase was initiated at a random intensity ranging between 0.2mA and 0.7mA, and between 0.7mA and 1.2mA, respectively. If the participant rated a stimulus >5, the intensity of the next step of the staircase would decrease by 0.1mA. If the participant rated a stimulus <5, it would increase by 0.1mA. If the participant rated a stimulus = 5, the intensity did not change. After both staircases were finished, the final values of both staircases were averaged, and that value was used as the stimulation intensity for the remainder of the experiment. Participants were instructed that the goal of this procedure was to ensure the electrical stimulation was unpleasant, but not extremely painful.

#### 2.2.2 Conditioning Phase

After the work-up procedure, participants were exposed to a conditioning phase in which they observed six different fractal stimuli (CSs) on screen (Figure 1A). CS conditions to which the stimuli were assigned were counterbalanced across participants. Two stimuli (CS-) were instructed to never be followed by an electrical stimulus (US). Two further stimuli were instructed to be followed by an electrical stimulus on some trials (CS+E), and the same instruction was given for the last two stimuli (CS+I). The key difference between CS+E and CS+I was that the former were indeed sometimes followed by an electrical stimulus (33% of the trials, chosen randomly), while the latter were not. Before the start of the conditioning phase, participants were asked to correctly identify each CS as either CS+ or CS-, and the experiment only started after they passed this test without errors.

**Figure 1.**
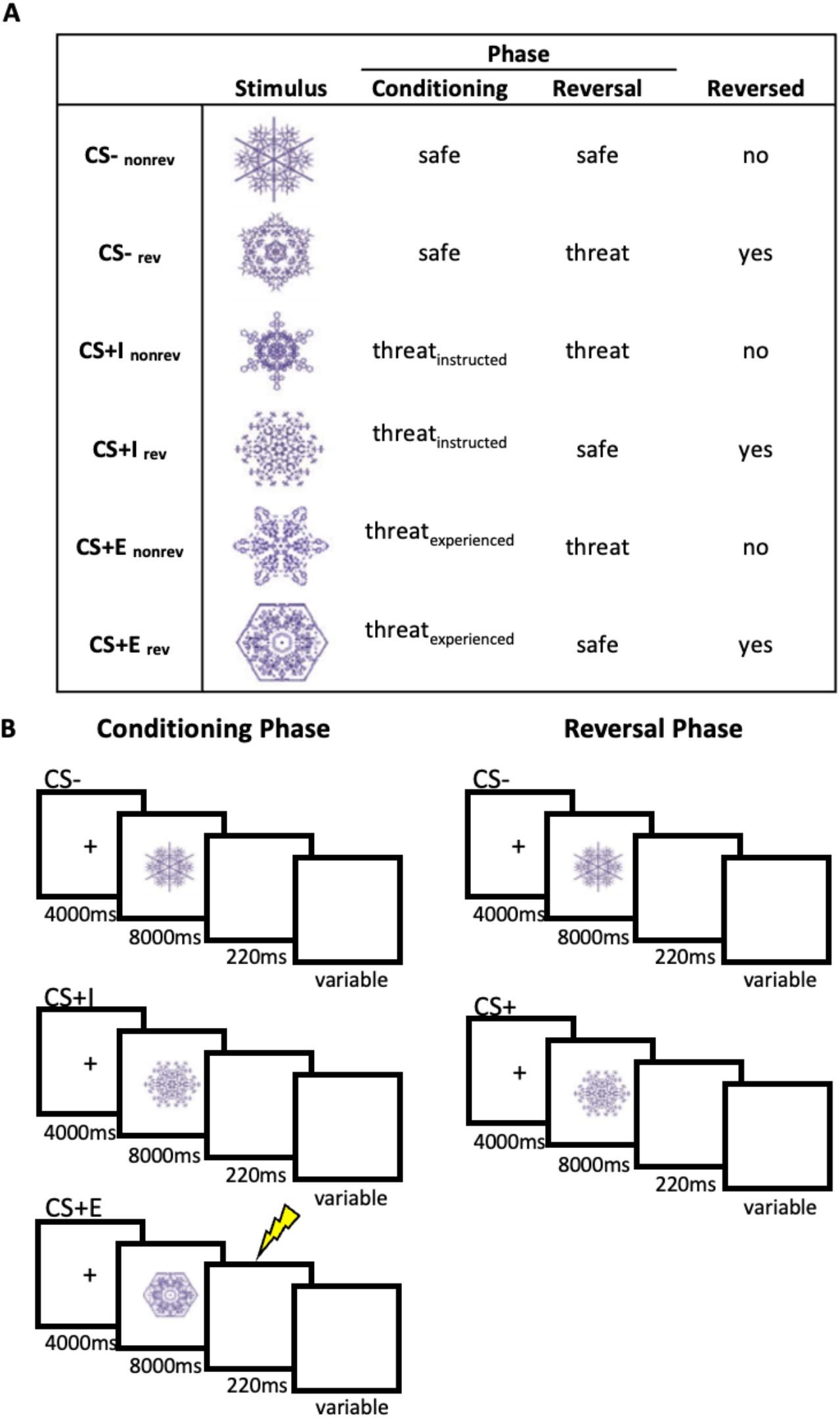
Stimuli and design. **A**. Conditions and stimuli used in this experiment. Stimulus to condition mappings were counterbalanced across participants. **B**. Trial structure in the conditioning and reversal phases, separately for each CS condition. The lightning bolt represents reinforcement, which was administered on 33% of the CS+E trials. In the reversal phase, stimuli were either instructed to be safe or threatening, but no stimuli were reinforced. CS-= safe conditioned stimulus, CS+I = instructed only conditioned stimulus, CS+E = instructed and experienced conditioned stimulus.

The conditioning phase consisted of a total of 36 trials (6 repetitions of each CS, Figure 1B). Each trial started with the presentation of a fixation cross (4000ms) centrally on screen. This was followed by the presentation of a CS on screen (8000ms). On reinforced trials (33% of CS+E trials), this was followed by an electrical stimulation, on all other trials a blank screen was presented for 220ms, which was the same duration as the electrical stimulation and was included to match all trials with respect to their duration. After a variable inter-trial-interval (13, 15, or 17s) the next trial started. Conditioning trials were presented in a random order, and randomization was performed in mini-blocks of 6 trials that contained each CS once. No direct repetitions of the same CS were allowed. Additionally, we ensured that CS condition was not correlated with either ITI duration or reversal condition, and that that the CS condition on a particular trial could not be predicted on the basis of the previous trial.

After 12 trials, the experiment was paused and a rating block was presented. Participants were asked to rate their self-reported CS fear and US expectancy for each of the CSs presented in the last 12 trials. Each CS was presented twice, once with the question “How fearful were you while seeing this stimulus?” (fear rating, 9-point Likert scale anchored at 1 “not at all”, 3 “rather not”, 5 “unsure”, 7 “somewhat”, 9 “strongly”), and once with the question “To which degree did you expect an electric shock while seeing this stimulus?” (US expectancy rating, 9-point Likert scale anchored at 1 “not at all”, 3 “a little”, 5 “average”, 7 “somewhat”, 9 “strongly”). Participants were instructed to base their ratings on the last time they saw each CS and responded in their own time. Once they answered each of the 12 rating items the experiment continued. After the next 12 trials, another rating block was presented, and after the last 12 trials the third and last rating block was presented, splitting up the conditioning phase into three blocks separated by rating items. Overall, the condition phase lasted about 18 minutes in total and consisted of a single fMRI run.

#### 2.2.3 Reversal Phase

After the conditioning phase, participants were instructed that some of the CSs would change their meaning in the next phase (Figure 1A). One CS-would remain a CS-(CS-_norev_), while the other could now be followed by an electrical stimulation (CS-_rev_), and two out of four CS+s would now be safe. Specifically, one CS+I would remain a CS+ (CS+I_norev_), while the other would not be followed by an electrical stimulus anymore (CS+I_rev_), and one CS+E would remain a CS+ (CS+E_norev_), while the other would not be followed by an electrical stimulation anymore (CS+E_rev_). Please note that participants were not instructed on this difference between CS+Es and CS+Is, they merely received threat / safety instructions for each CS. Thus, the overall design of this study was a 3 (CS conditions: CS-, CS+E, CS+I) x reversal (reversed, non-reversed) x time (conditioning phase, reversal phase). Similar to the conditioning phase, the reversal phase only started after the participant correctly indicated the updated CS-US association for each CS. It was stressed that the second phase would be similar to the first phase, with some CSs to be followed by a US on a portion of the trials, and other CSs never followed by a US. However, similar to the study by Braem and colleagues (Braem et al., 2017), no electrical stimulation was applied in the reversal phase (Figure 1B).

The overall structure of the second run (reversal phase) was identical to the conditioning phase, with updated CS contingencies. Half of the CSs reversed, while the other retained their original association, and no electrical stimulation was administered in the reversal phase. After each block of 12 trials, participants were asked to rate their fear and US expectancy for each CS. The reversal phase lasted about 18 minutes, and consisted of a single fMRI run.

#### 2.2.4 Debriefing

After the reversal phase, participants left the MR scanner, were debriefed, and filled in the state trait anxiety index (STAI, trait version, Spielberger et al., 1999). Within the context of this paper, we did not further investigate STAI results, and they are not reported here. All participants were further asked to which degree they believed the instructions, at the time they were given. Almost all participants believed the instructions, only two indicated the believability to be somewhat low.

### 2.3 Skin Conductance Response Acquisition

For each subject, skin conductance responses (SCR) were collected using Biopac hardware (EDA100C-MRI, PPG100C) and standard disposable Ag/AgCl electrodes attached to the thenar and hypothenar eminences of the non-dominant (left) hand. The signal was measured using the ACQKnowledge softare and digitized at 2000Hz. For the SCR analysis, trials in which subjects were presented a US were excluded, as the electrical shocks might bias the SCRs. Data were further analyzed using Matlab (R2014b 8.4.0 150421, Mathworks, RRID: SCR_001622). They were first smoothed using a Gaussian kernel, and SCRs were calculated by subtracting the mean value of a baseline time period (6-4s before CS onset) from the maximum amplitude within a 1-7s interval after the CS onset. Any values smaller than 0.02, including negative values, were scored as zero. Then, SCRs were range-corrected (Lykken and Venables, 1971) and square root transformed to normalize the data. This analysis procedure has been used successfully previously (Mertens and De Houwer, 2016). Unexpectedly, we were unable to detect any SCR signal above the baseline in the relevant response window (1-7s after CS onset) for any condition, and thus did not analyze SCRs any further.

### 2.4 Behavioral data analysis

#### 2.4.1 Manipulation check

In order to test whether participants understood the difference between threatening and safe CSs (CS-vs CS+E), we tested whether US expectancy ratings differed between these conditions, and whether they reversed in line with the verbal instructions. For this purpose, we performed a 3-factorial Bayesian ANOVA with the within-subject factors CS (CS-, CS+E), reversal (reversed, non-reversed), phase (conditioning, reversal), adding subjects as a random factor. All ANOVAs were computed in R (Rstudio, v1.1.456, RRID: SCR_000432) using the *BayesFactor* package. Results are reported in terms of Bayes factors (BF10, default inverse-chi-square prior, scaling factor = 0.5). Following previous research (e.g., Andraszewicz et al., 2015; Mertens and De Houwer, 2016), we considered BFs between 0.33 and 1 as anecdotal evidence, BFs between 0.1 and 0.33 as moderate evidence, and BFs smaller than 0.1 as strong evidence for the null hypothesis. BFs between 1 and 3 were considered as anecdotal evidence, BFs between 3 and 10 as moderate evidence, and BFs larger than 10 as strong evidence for an alternative hypothesis.

We tested for a main effect of threat, expecting higher US expectancy ratings for threatening (CS+E) than for safe (CS-) stimuli. We also tested whether reversal instructions increased ratings for reversed CS-, and decreased ratings for reversed CS+E, which should be seen as a 3-way interaction of CS, reversal, and phase (see Mertens and De Houwer, 2016, for more information on this analysis logic in a highly similar design). The same tests were conducted on the fear ratings.

#### 2.4.2 Experience effects

In our main analysis, we tested whether experiencing a US affected fear reversal, as measured using US expectancy ratings. For this purpose, we computed a 3-way Bayesian ANOVA again, using the factors CS (CS+I, CS+E), reversal (reversed, non-reversed), phase (conditioning, reversal), and modelling subjects as a random effect. By comparing CS+I and CS+E here, we were able to test whether reversals of CS-US contingencies across phases differed between CS+I and CS+E, in which case we should find evidence for a 3-way interaction of CS, reversal, and phase. In order to test whether experience effects change over the course of the experiment, we first split each phase into three separate blocks (separated by rating items). We then estimated an additional Bayesian ANOVA, using the same three factors listed above, only adding block as an additional factor.

In order to investigate the effects of prior CS-US experience after verbal reversal instructions, we performed a number of tests on the reversal phase only. First, we tested whether expectancy ratings differed between CS-_rev_, CS+I_norev_, and CS+E_norev_ using a one-factorial Bayesian ANOVA, modelling subjects as a random factor. All of these CSs were threatening during the reversal phase, and only differed in their history during the conditioning phase. While CS-_rev_ only just became threatening, CS+E_norev_ was followed by USs and remained threatening, and CS+I_norev_ was not followed by USs and remained threatening. If learning history carried over from the conditioning to the reversal phase, we should see stronger responses to CS+E_norev_ than to either CS-_rev_ and CS+I_norev_. We used a similar approach to compare the three safe stimuli in the reversal phase, CS-_norev_, CS+I_rev_, and CS+E_rev_. These stimuli only differed in their learning history in the conditioning phase, and if this history carried over to the reversal phase, we would expect the see higher US expectancy for CS+E_rev_ than for CS+I_rev_ and CS-_norev_. Again, the same analyses were then performed on fear ratings as well. In line with Mertens and De Houwer (2016), we expected no additional effects of having experienced CS-US pairings.

### 2.5 fMRI data acquisition

fMRI data was collected using a 3T Magnetom Trio MRI scanner system (Siemens Medical Systems, Erlangen, Germany), with a standard thirty-two-channel radio-frequency head coil. A 3D high-resolution anatomical image of the whole brain was acquired for co-registration and normalization of the functional images, using a T1-weighted MPRAGE sequence (TR = 2250 ms, TE = 4.18 ms, TI = 900 ms, acquisition matrix = 256 × 256, FOV = 256 mm, flip angle = 9°, voxel size = 1 × 1 × 1 mm). Furthermore, a field map was acquired for each participant, in order to correct for magnetic field inhomogeneities (TR = 400 ms, TE_1_ = 5.19 ms, TE_2_ = 7.65 ms, image matrix = 64 × 64, FOV = 192 mm, flip angle = 60°, slice thickness = 3 mm, voxel size = 3 × 3 × 3 mm, distance factor = 20%, 33 slices). Whole brain functional images were collected using a T2*-weighted EPI sequence (TR = 2000 ms, TE = 30 ms, image matrix = 64 × 64, FOV = 192 mm, flip angle = 78°, slice thickness = 3 mm, voxel size = 3 × 3 × 3 x mm, distance factor = 20%, 33 slices). Slices were orientated along the AC-PC line for each participant.

### 2.6 fMRI data analysis

#### 2.6.1 Preprocessing

fMRI data was analyzed using Matlab (The MathWorks, version R2014b 8.4.0 150241, RRID: SCR_001622) and the SPM12 toolbox (http://www.fil.ion.ucl.ac.uk/spm/software/spm12/, version 6906, RRID: SCR_007037). Before the analysis, we discarded the first three acquired volumes of each run. Functional data was subsequently unwarped, realigned, and slice-time corrected. The preprocessed data was then screened for possible scanner-related artifacts, using the Artifact Detection Tool (http://www.nitrc.org/projects/artifact_detect/, ART version 2011-07, RRID: SCR_005994). ART automatically detects and marks outlier volumes based on the global mean brain activation, and movement parameters (z-threshold=9, movement threshold=2). We identified outlier volumes in 17 participants (mean number of identified volumes = 2.7, min = 2, max = 28). Variance attributable to these artifacts was removed by explicitly modelling the affected volumes in the respective first-level general linear models (GLMs).

#### 2.6.2 First-level GLM estimation

After preprocessing, a GLM (Friston et al., 1994) was estimated using unsmoothed, non-normalized data, separately for each participant (GLM_main_). This allowed us to perform all representational similarity analyses in native space for each participant. Each combination of CS type (CS-, CS+I, CS+E), reversal (reversal, no reversal), and phase (conditioning, reversal) was modelled using a separate regressor. This resulted in 12 regressors of interest in this GLM. Each regressors was modelled as a boxcar locked to the onset of the CS presentation (duration = 8 s), and was convolved with a canonical HRF basis-function, an approach used successfully before (Braem et al., 2017). Regressors of non-interest included the onset of each US, one regressor per identified outlier/artifact volume (as suggested in the ART documentation), and 6 movement regressors.

Next, we estimated a second GLM for each subject (GLM_block_). This GLM was identical to GLM_main_, only that we added block as an additional factor. As outlined above, each phase consisted of three blocks, separated by rating items. Adding block as a factor allowed us to investigate the temporal evolution of any observed effects, and allowed us to restrict analyses to e.g. only the first block of each phase, where instruction effects should be strongest. Overall, this GLM included 36 regressors of interest (CS x reversal x phase x block), and regressors of non-interest were identical to the previous GLM.

#### 2.6.3 ROI definition

Our main analyses were performed within several a-priori defined regions of interest (Figure 2) that have been found to be involved in fear conditioning (Visser et al., 2013; Fullana et al., 2015). Specifically, we included ROIs for the bilateral anterior cingulate cortex (ACC), bilateral insula (INS), bilateral ventral striatum (VS), bilateral thalamus (TH), bilateral ventromedial PFC (vmPFC), bilateral superior frontal gyrus (SFG), left amygdala (lAMY), and right amygdala (rAMY)). These ROIs were obtained from the Harvard-Oxford cortical and subcortical structural atlas (Harvard Center for Morphometric Analysis), using a probability threshold of 25%. Additionally, we constructed a spherical ROI (radius = 10mm) in the right dorso-lateral PFC (dlPFC), using coordinates from a previous paper ([39 20 31], Demanet et al., 2016, as also used in Bourguignon et al., 2018) and the WFUpickatlas toolbox (version 3.0.5, RRID: SCR_007378, Maldjian et al., 2003). This region has been shown to be uniquely involved in instruction implementation of task rules, and could therefore also be involved in implementing the reversal instructions in our dataset. All ROIs were then projected into native space, separately for each participant, using the inverse normalization field estimated during preprocessing.

**Figure 2:**
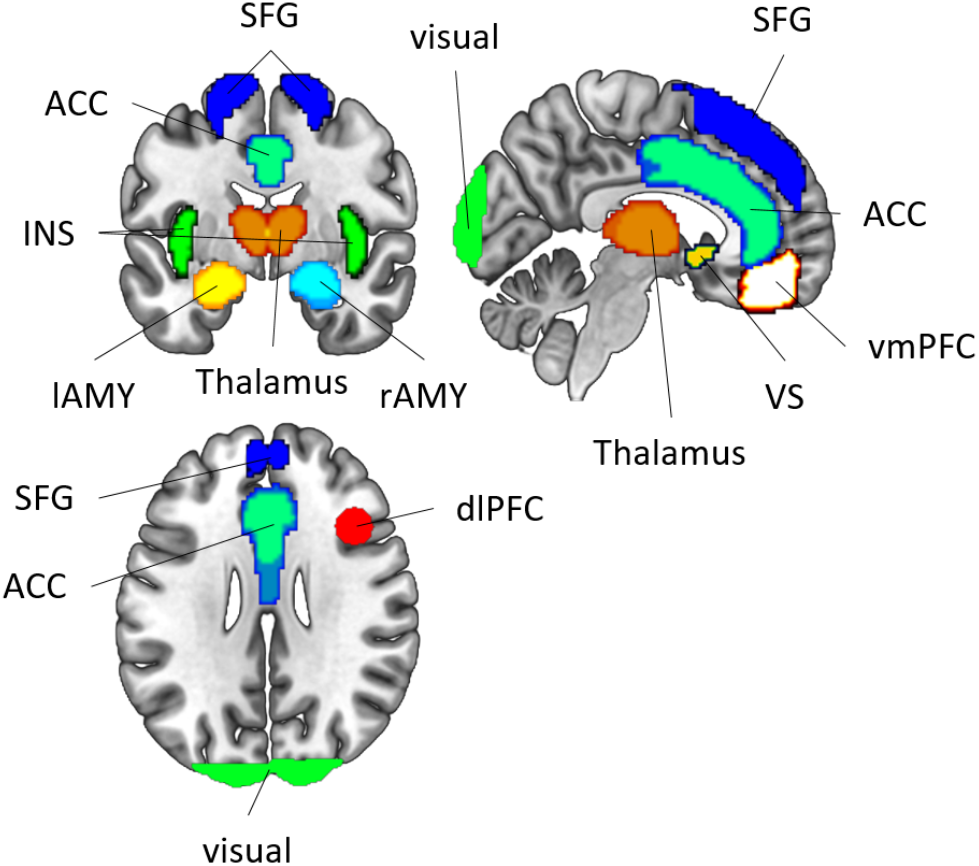
Regions-of-interest. Anterior cingulate cortex (ACC), superior frontal gyrus (SFG), ventromedial prefrontal cortex (vmPFC), dorsolateral prefrontal cortex (dlPFC), ventral striatum (VS), thalamus, left amygdala (lAMY), right amygdala (rAMY), insula (INS).

#### 2.6.4 Representational Similarity Estimation

We then used the beta estimates from the GLM_main_ to perform representational similarity analyses, RSA (Kriegeskorte et al., 2008). This allowed us to measure the representational distances between individual CSs, and thus test whether neural CS representations were more or less similar to each other depending on threat and / or prior CS-US experience. For each ROI, we first extracted beta values for each of the 12 conditions in this experiment, i.e. each combination of CS x reversal x phase. Since we were unable to implement a leave-one-run-out cross-validation procedure with the current design, we instead z-scored the data across phases in order to avoid confounds related to global signal differences between the different runs of the experiment. Please note that all RSAs were performed in native space for each participant. Then, we performed multivariate noise normalization (Walther et al., 2016), after which we calculated Pearson correlation coefficients for each pairwise comparison of the 12 conditions. This resulted in a 12×12 correlation matrix (representational similarity matrix, RSM). Before running statistical tests on these correlations, they were first Fisher-z transformed. At the group level, correlations were assessed using either Bayesian t-tests (Cauchy prior, scaling factor = 0.71) or Bayesian ANOVAs (inverse-chi-square prior, scaling factor = 0.5).

#### 2.6.5 Exploratory analyses

##### Model based RSA of expectancy and fear ratings

There is an ongoing debate about the differences and similarities between expectancy and fear ratings (Mertens and De Houwer, 2016; Mertens et al., 2018), and in order to explore this issue in more detail, we also performed an additional exploratory analysis using model-based searchlight RSA (Nili et al., 2014). First, we computed two theoretical model-RSMs from behavioral US expectancy ratings and fear ratings, respectively, by computing pairwise correlations between these ratings for all CSs. Then, we used a searchlight approach (radius = 3 voxels, Etzel et al., 2013) to determine in which brain regions neural RSMs matched the theoretical predictions. In each searchlight, we computed the partial correlation between one model RSM and the data RSM, while controlling for the influence of the other model, e.g. r(US expectancy model, data) while controlling for the fear model. This resulted in a whole-brain similarity map, showing which brain regions share unique variance with US expectancy ratings while controlling for fear ratings, and vice versa. Maps were then smoothed (6mm FWHM) and normalized (MNI-template, as implemented in SPM12). On the group level, we performed two separate GLMs, one for each model, testing where its unique shared variance with the data RSM was larger than 0, and results were corrected for multiple comparison using a voxel threshold of p < 0.05 (FWE corrected). This analysis allowed us to directly test which brain regions were specifically associated with either expectancy or fear ratings, potentially leading to new hypotheses that can be tested in future research.

## 3 Results

### 3.1 Behavioral results

#### 3.1.1 Manipulation check

We first contrasted safe (CS-) and experienced (CS+E) stimuli to test for an effect of threat. For US expectancy ratings (Figure 3a), we found strong evidence for a main effect of threat (BF10 > 150), as well as strong evidence for a 3-way interaction of CS-type (CS-vs CS+E), reversal, and phase (BF10 > 150). For fear ratings (Figure 3b), we found highly similar results: a main effect of threat (BF10 > 150), and a 3-way interaction (BF10 > 150). These results demonstrated that participants clearly perceived threatening and safe stimuli differently, and that reversal instructions differentially affected CS- and CS+E stimuli, increasing ratings for reversed CS- and decreasing ratings for reversed CS+E.

**Figure 3:**
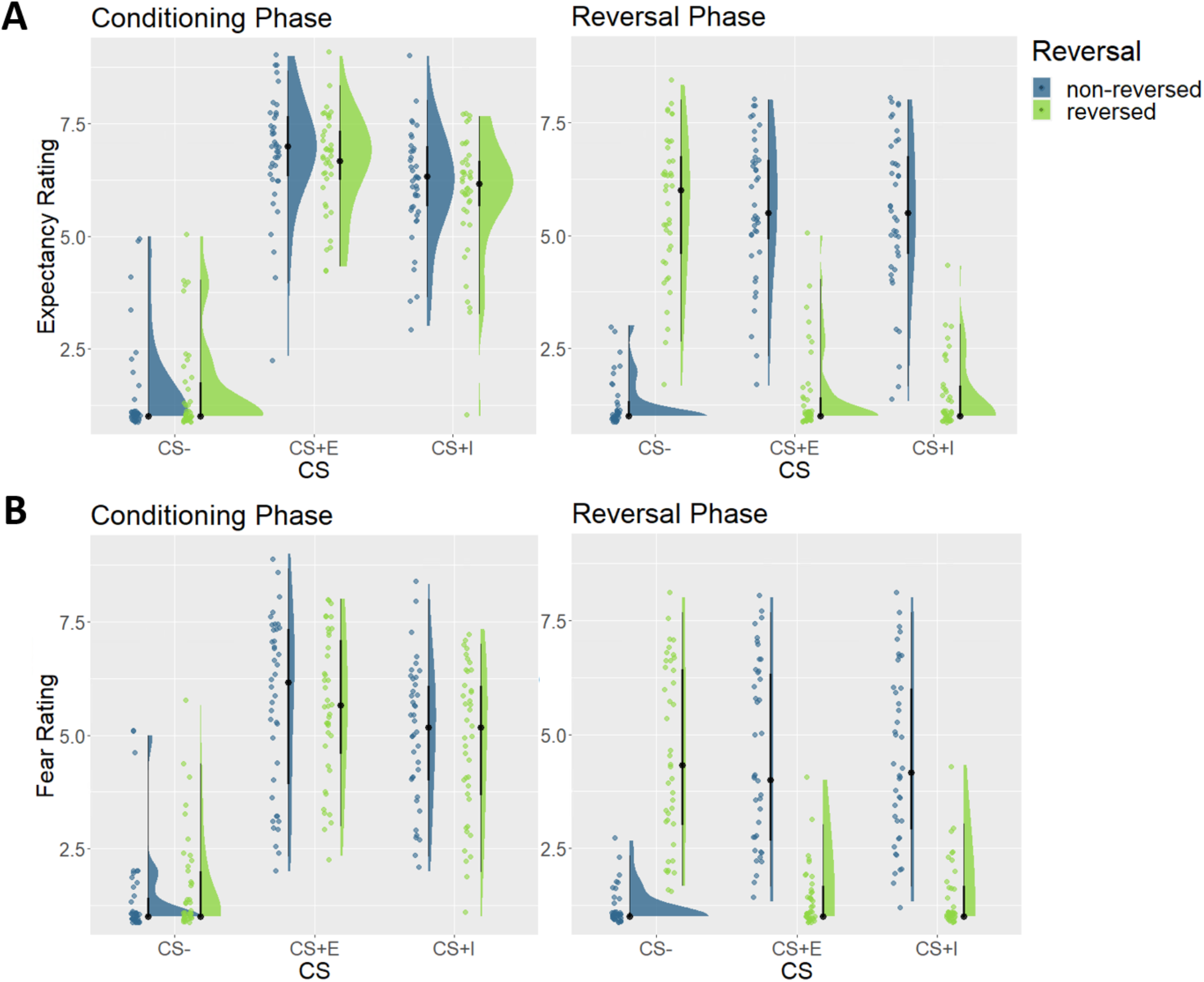
Expectancy and fear ratings. **A**. US expectancy ratings for the conditioning phase (left), and the reversal phase (right). **B**. Fear ratings for the conditioning phase (left), and the reversal phase (right). Ratings were made on a 9-point Likert scale. CS-= safe conditioned stimulus, CS+I = merely instructed conditioned stimulus, CS+E = instructed and experienced conditioned stimulus.

#### 3.1.2 Experience effects

To test for the effect of CS-US experience, we then contrasted CS+I with CS+E. For US expectancy ratings, we found anecdotal evidence against a main effect of experience, suggesting that experienced stimuli showed similar ratings as merely instructed stimuli, BF10 = 0.38. Furthermore, we found moderate evidence against a 3-way interaction of CS-type (CS+E vs CS+I), reversal, and phase, BF10 = 0.23. This suggests that reversal instructions affected experienced and merely instructed stimuli to an equal degree. We then assessed expectancy ratings in the reversal phase in more detail. If the learning history affected behavior, US expectancy ratings should be higher for safe stimuli that were threatening in the past (CS+E_rev_), as compared to safe stimuli that were safe throughout the experiment (CS-_norev_). However, we found evidence against this hypothesis, BF10 = 0.28. Conversely, one would also expect threatening stimuli that were safe in the past (CS-_rev_) to show lower ratings, as compare to threatening stimuli that were threatening throughout the experiment (CS+E_norev_). We again found evidence against this hypothesis, BF10 = 0.11.

One potential reason why we found no differences between CS+E and CS+I stimuli in the reversal phase could be due the fact that US expectancies decayed over the course of the reversal phase. Indeed, we found strong evidence for block effects in the reversal phase, BF10 > 150, demonstrating that US expectancy ratings were higher in the beginning than in the end of the reversal phase. We repeated the analysis reported above, now only using data from the first block of each phase, where instructions should have the strongest impact. Even here, we found evidence against differences between CS+E and CS+I, BF10s < 0.29. Thus, in line with Mertens and De Houwer (2016), we found evidence against an effect of CS-US experience on expectancy ratings in the reversal phase.

For fear ratings, we found that ratings were overall higher for experienced (CS+E), as compared to merely instructed stimuli (CS+I), BF10 = 3.43, which was mainly driven by differences in the conditioning phase. We again found moderate evidence against a 3-way interaction however, BF10 = 0.18, showing that reversal instructions had comparable effects on experienced and merely instructed stimuli. In the reversal phase, we found no evidence for an effect of learning history on ratings of threatening (BF10 = 0.09) and inconclusive evidence for a similar effect on safe stimuli (BF10 = 1.54). Additional exploratory analyses demonstrated that fear ratings also showed extinction effects in the reversal phase (BF10 > 150), yet restricting the analysis only to the first rating block still yielded evidence against differences between CS+E and CS+I, BF10s < 0.31. Overall we replicated Mertens and De Houwer (2016), finding evidence against an effect of CS-US experience on fear ratings in the reversal phase.

Note that we found evidence for a main effect of CS-US experience in the fear ratings, but not in the US expectancy ratings, driven by differences in the conditioning phase. In order to test whether this difference was robust, we repeated the ANOVA described above (factors: CS (CS+I, CS+E), reversal, phase), adding an additional factor item type (US expectancy, fear). We then assessed whether we found evidence for a CS x item type interaction, which would indicate that the main effect of CS (CS+E vs CS+I) was stronger for fear than for expectancy ratings. We found moderate evidence against this hypothesis, BF10 = 0.13. Thus, differences between US expectancy and fear ratings should be interpreted with caution.

### 3.2 fMRI results

#### 3.2.1 Manipulation check

##### Conditioning phase

As a first manipulation check, we assessed whether the experience of CS-US pairings affected voxel pattern responses to the different CSs in the conditioning phase (Figure 4). If a region were responsive to instructed threat, we would expect the pattern similarity between threatening (CS+E, CS+I) and safe CSs (CS-) to be lower than between two threatening CSs (CS+E, CS+I). We thus estimated the representational similarities between each CS type in the conditioning phase, collapsing across reversal conditions here since this factor became relevant only after the reversal instructions. We then performed paired Bayesian t-tests to test whether r(CS+E, CS+I) was higher than either r(CS+E, CS-) and r(CS+I, CS-). We found strong evidence for this effect in every ROI included here, all BF10s > 150 (Figure 5), suggesting that each fear-related brain region encoded instructed threat.

**Figure 4:**
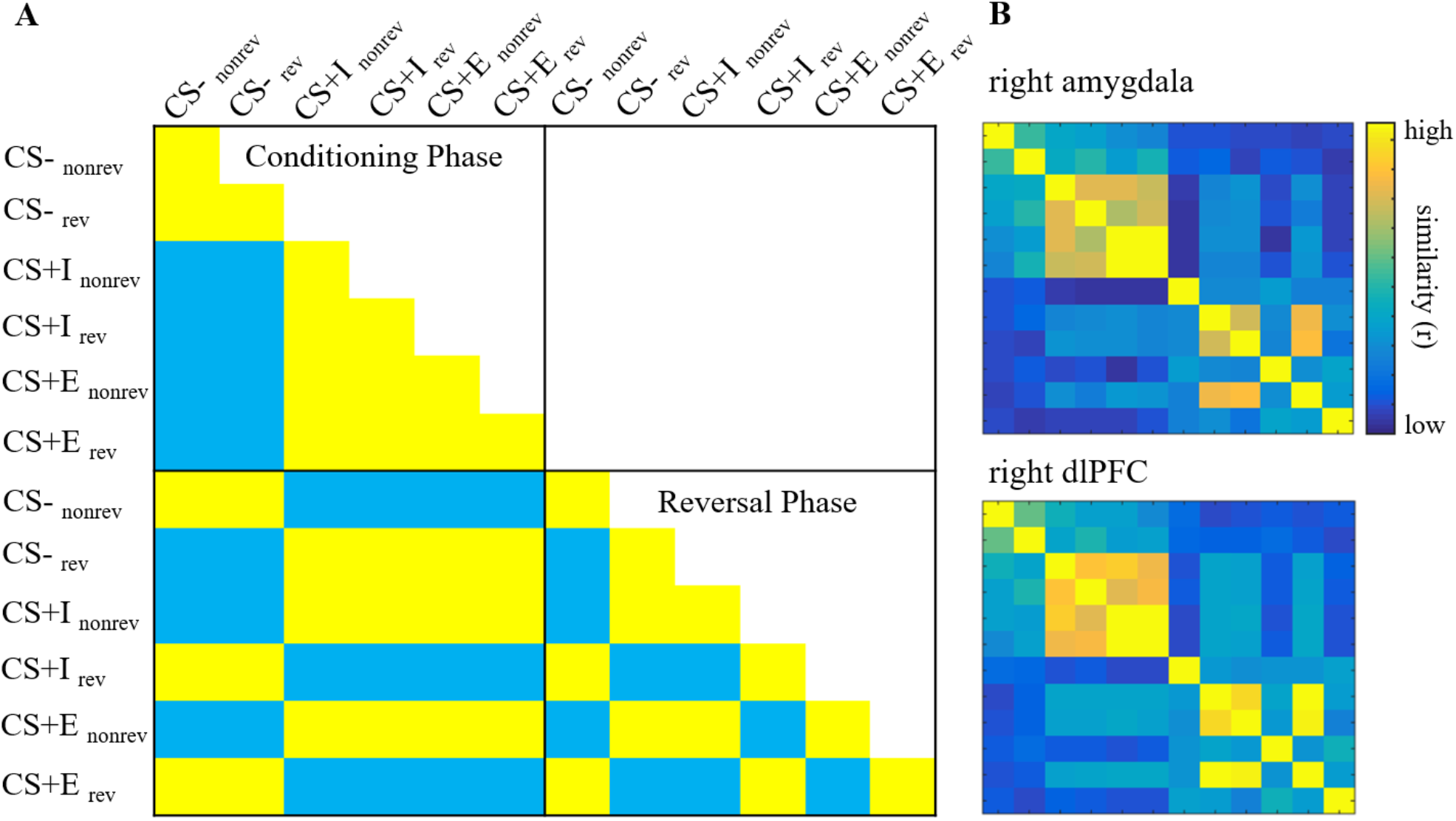
Representational similarity matrices (RSM). **A**. Theoretical threat model RSM. Each cell depicts the expected pairwise correlations for each condition if a region coded for the presence of threat. Conditions: CS type (CS-, CS+I, CS+E), reversal (rev = reversed, nonrev = not reversed), phase (Conditioning Phase, Reversal Phase). **B**. RSMs extracted from two example regions of interest, the right amygdala, and the right dorsolateral prefrontal cortex (dlPFC).

**Figure 5.**
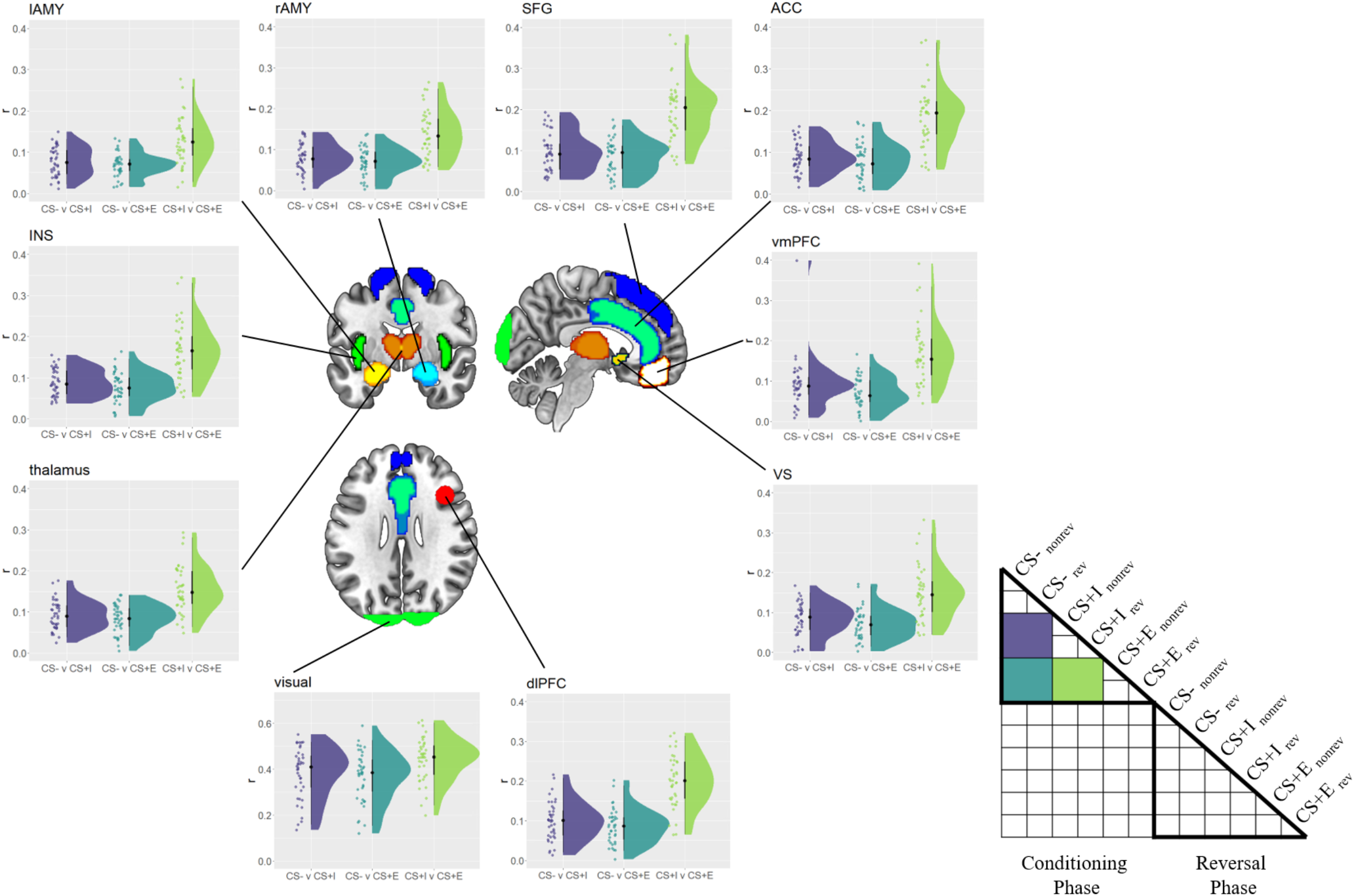
Between-CS similarity in the conditioning phase. Similarity (r) between safe (CS-), merely instructed (CS+I) and instructed + experienced (CS+E) CSs in the conditioning phase, separately for each ROI. Raincloud plots depict the raw data as dots on the left side and the data distribution on the right side. The schematic RSM denotes which pairwise representational similarity measures were used in this analysis.

Next, we tested whether ROIs showed more consistent voxel pattern responses to threatening stimuli (CS+E, CS+I) than to safe stimuli (CS-) in the conditioning phase. There is some prior evidence that threat not only changes the overall activity level in fear-related brain regions, but also induces more consistent voxel pattern responses to threatening CSs (Visser et al., 2011; Braem et al., 2017), similar to emotional stimuli (Riberto et al., 2022). Moreover, and different to the study by Braem and colleagues (2017), here we could test whether the ROIs provide more consistent voxel patterns within CS-type while controlling for low-level visual features. Namely, we tested this hypothesis by first computing the correlation between to be reversed and not to be reversed CS-s, r(CS-norev, CS-rev). Please note that at this time in the experiment, the meaning of these two stimuli is identical to all participants, they solely differ in, and therefore control for, their visual appearance. The same approach was used to estimate consistency of CS+E and CS+I. Following previous findings, we expected CS-s to show lower coding consistencies than CS+s, which we tested using Bayesian paired t-tests. We found strong evidence for more consistent coding of CS+E, as compared to CS-in every ROI (BF10s > 150, Figure 6). Furthermore, we found evidence for more consistent coding of CS+Is, as compared to CS-s, BF10s > 21.4, in almost all ROIs. The only exception here was the left Amygdala, in which we found no evidence for (or against) differences between CS+I and CS-, BF10 = 0.98. To explore this finding further, we directly compared consistencies in the left and right amygdala, to see if there is evidence for a hemispheric difference. Running an ANOVA using the factors CS (CS+I, CS-), and hemisphere (left, right), we found no evidence for (or against) either a main effect of hemisphere, BF10 = 1.00, or an interaction of hemisphere and CS, BF10 = 0.71. Thus, differences between left and right amygdala should be interpreted with caution.

**Figure 6.**
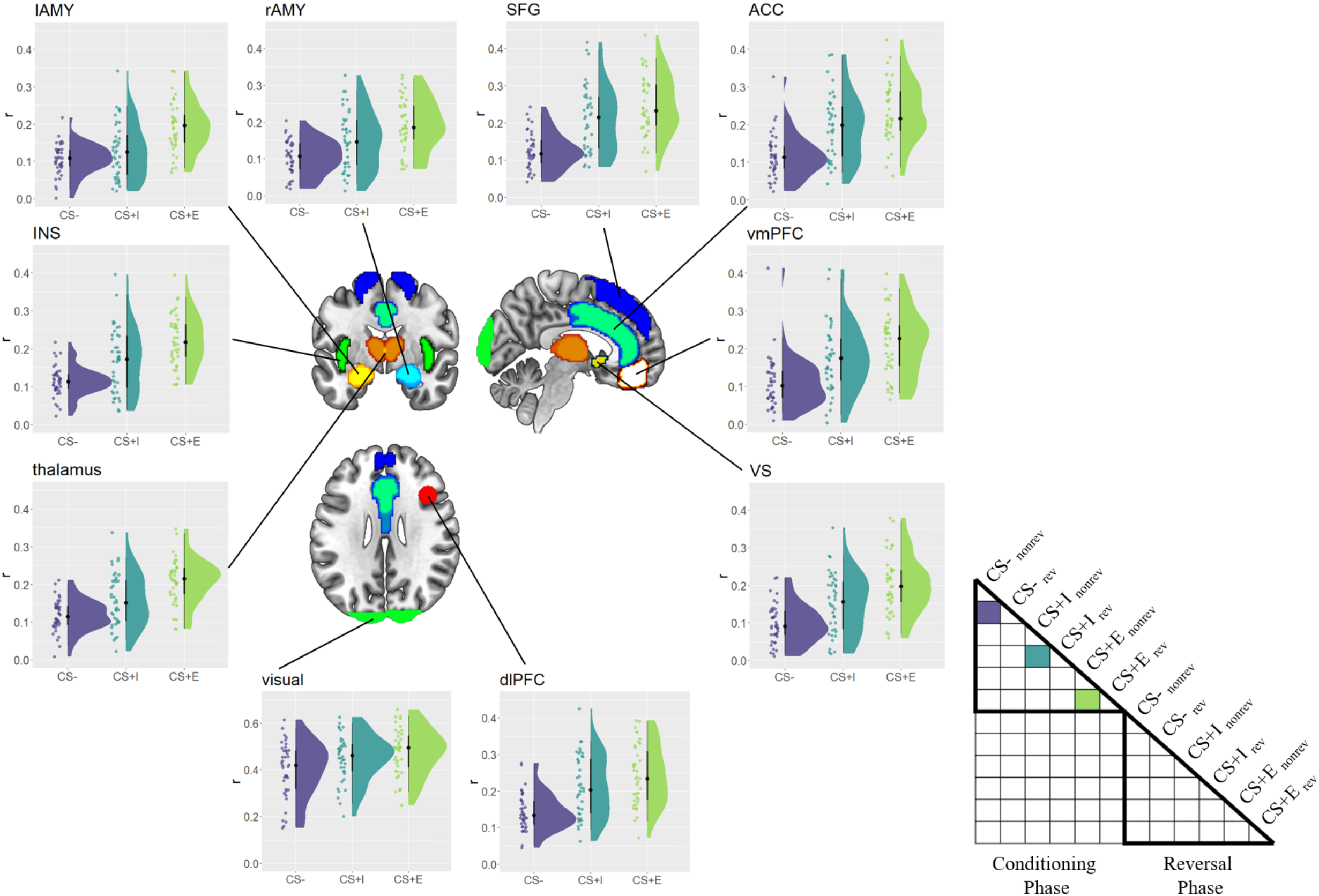
Within-CS consistency in the conditioning phase. Consistency (r) with which safe (CS-), merely instructed (CS+I) and instructed + experienced (CS+E) CSs were represented, separately for each ROI. Raincloud plots depict the raw data as dots on the left side and the data distribution on the right side. The schematic RSM denotes which pairwise representational similarity measures were used in this analysis.

##### Reversal phase

In the reversal phase, no reinforcements were given, making CS+I and CS+E identical with respect to the current, albeit not past, threat experiences. Therefore, we were unable to repeat the manipulation check reported above, which relied on comparing r(CS+E, CS+I) with r(CS+E, CS-). We were, however, able to investigate the consistency of the voxel pattern response in a similar manner to the conditioning phase. Since no US was presented, we first collapsed across the CS+E and CS+I conditions, then separately computed coding consistency for safe CSs (CS-non-reversed, CS+I reversed, CS+E reversed) and threatening CSs (CS-reversed, CS+I non-reversed, CS+E non-reversed), respectively. We found coding consistency to be lower for safe CSs than for threatening CSs in the reversal phase in each ROI, all BF10s > 150. This mirrors findings from the conditioning phase, even though no reinforcements were given here.

#### 3.2.2 Experience effects

##### Conditioning phase

In order to test whether CS-US experience comes with a unique neural trace, we first investigated BOLD responses in the conditioning phase. We reasoned that if experience had an effect above and beyond verbal instructions, different CSs that were actually paired with a US might share more variance with each other than with threatening CSs that were merely instructed. To test this, we first extracted the correlation between CS+E reversed and CS+E non-reversed during the conditioning phase, which captures any shared variance between two CSs with identical meaning during the conditioning phase but different visual features. In order to estimate the shared variance between different threatening CSs, we then computed and averaged the following correlations: r(CS+E reversed, CS+I reversed), r(CS+E reversed, CS+I non-reversed), r(CS+E non-reversed, CS+I reversed), and r(CS+E non-reversed, CS+I non-reversed). This measure served as our baseline and captured any shared variance between different threatening CSs, with different visual features and different CS-US experience, and therefore captures a general threat signal that is independent from CS-US experience. By computing the difference r(CS+E reversed, CS+E non-reversed) – baseline, we can then test whether two experienced CSs share more variance with each other than they do with other threatening CSs which were not experienced. We tested this hypothesis using Bayesian one-sided t-tests against zero, and this served as one key measure of experience effects on neural coding in the conditioning phase. We found strong evidence for such an experience effect in the conditioning phase in each ROI, BF10 > 150 (Figure 7). Additionally, we explored whether CS+Is also showed a similar effect. We had no strong a priori hypotheses about this test, and we found an effect that was indistinguishable from 0 in each ROI, BF10s < 0.37. These results indicate that during conditioning, different experienced CS+s share more variance with each other than they do with other, merely instructed CS+s, indicating experience effects that go beyond verbal threat instructions.

**Figure 7:**
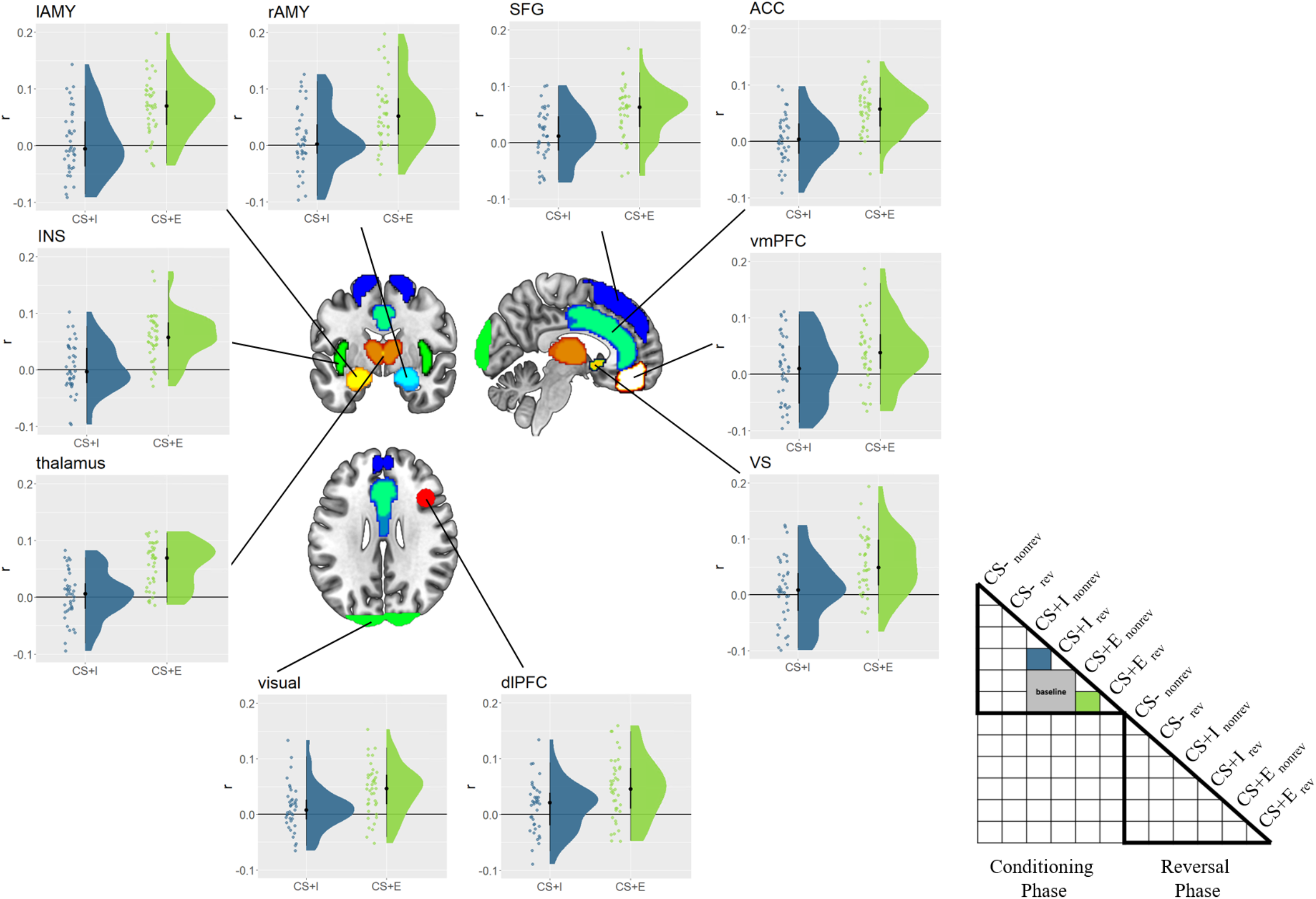
Experience effects in the conditioning phase. Similarity values for merely instructed (CS+I) and instructed + experienced (CS+E) CSs are depicted for each ROI, with the baseline subtracted. Raincloud plots depict the raw data as dots on the left side and the data distribution on the right side. The schematic RSM denotes which pairwise representational similarity measures were used in this analysis, including the baseline conditions (grey).

##### Reversal phase

Next, we evaluated our main research question: Whether CS-US experience has an effect on neural responses above and beyond verbal instruction effects after reversal. Our behavioral results and prior research showed that verbal instructions alone can fully reverse and explain effects on US expectancy and psychophysiological measures (Mertens and De Houwer, 2016), with no unique effect attributable to CS-US experience. To test whether there was still a trace of CS-US experience in the neural voxel pattern responses, we first extracted the correlation between CS+E reversed (now safe) and CS+E non-reversed (still threatening), and between CS+E reversed (now safe) and CS+I non-reversed (still threatening). The two threatening stimuli included here solely differ in their past learning experience. If there was a lingering representation of threat due to past CS-US experience, we would expect r(CS+E reversed, CS+E non-reversed) to be larger than r(CS+E reversed, CS+I non-reversed). This comparison controls for the mere presence of threat, and instead aims to isolate a unique effect of prior experienced threat only, making this a highly conservative analysis approach. We tested this hypothesis using Bayesian paired t-tests, r(CS+E reversed, CS+E non-reversed) vs r(CS+E reversed, CS+I non-reversed), separately for each ROI, and expected to find evidence for prior CS-US experience only in the right amygdala. Unexpectedly, we found evidence for prior CS-US experience in all brain regions assessed here, BF10s > 3.25, except the vmPFC, BF10 = 1.12, and right amygdala, BF10 = 0.44 (Figure 8). Thus, counter to our initial expectations, most threat-related brain regions showed evidence for lingering effects of prior CS-US experience, even after verbal reversal instructions.

**Figure 8:**
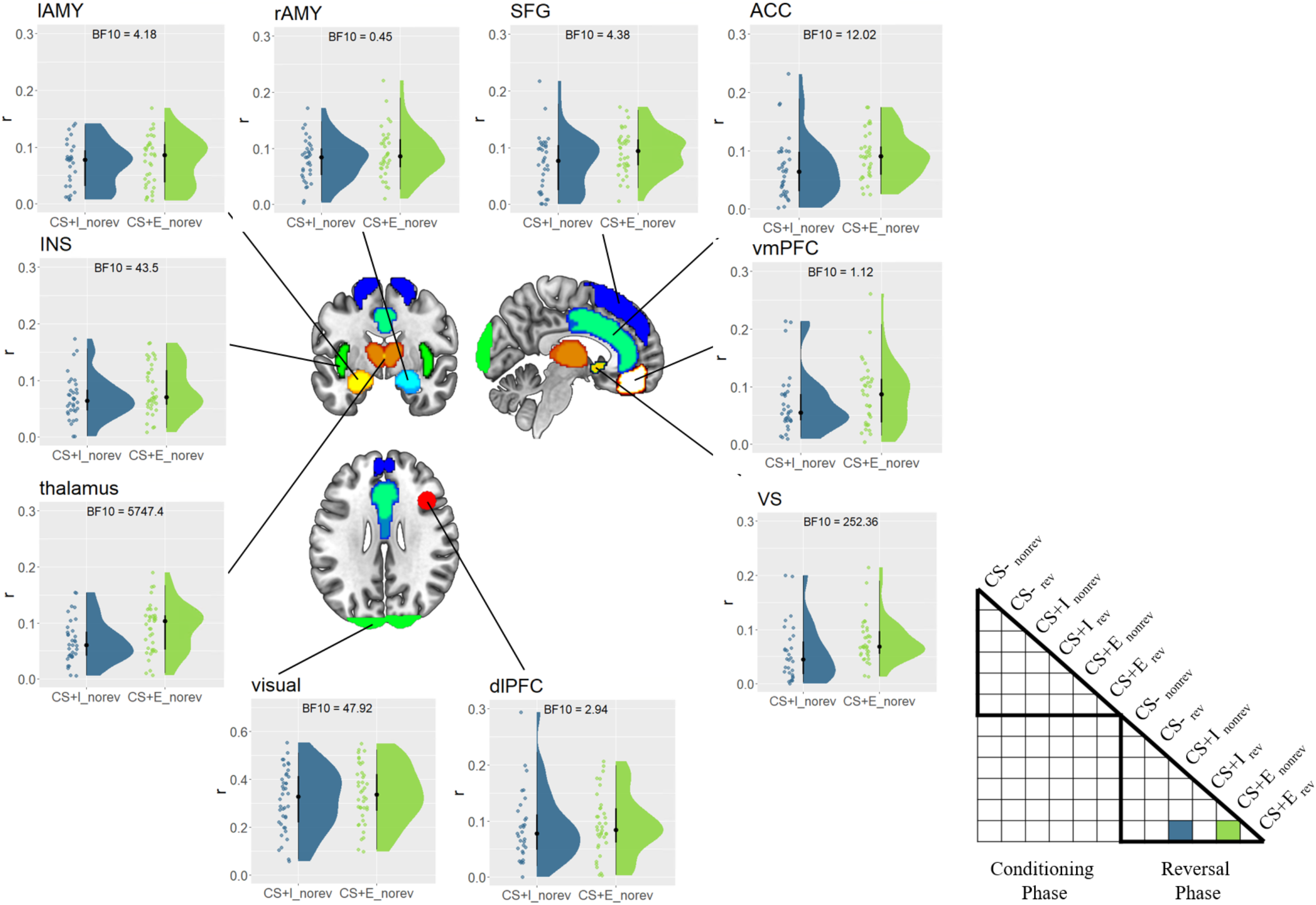
Effects of prior CS-US experience. Similarity between CS+E reversed (safe) and CS+I non-reversed / CS+E non-reversed (threatening) in the reversal phase, separately for each ROI. Higher similarities for CS+E reversed / non-reversed indicate a Pavlovian trace. Raincloud plots depict the raw data as dots on the left side and the data distribution on the right side. The schematic RSM denotes which pairwise representational similarity measures were used in this analysis.

In an additional post-hoc test, we assessed whether these effects decreased over time. We reasoned that even if many fear-related brain regions showed lingering effects of prior CS-US experience, they might still differ in their duration, i.e. in some regions effects might vanish over time, while in others they might persist. We first used GLM_block_ to calculate r(CS+E reversed, CS+E non-reversed) vs r(CS+E reversed, CS+I non-reversed), separately for each of the three blocks in the reversal phase. We then entered these values into a one-factorial Bayesian ANOVA (block), separately for each ROI. We found evidence against any differences between blocks in all ROIs, BF10s < 0.29, except the rAMY, where evidence remained inconclusive, BF10 = 0.46. Thus, effects of prior CS-US experience seem to be unaffected by presenting the CS without the US and seem to persist across time.

#### 3.2.3 Exploratory analyses

##### Model-based RSA of expectancy and fear ratings

In order to explore the relation of neural voxel pattern responses and the behavioral ratings in more detail, we also performed an additional exploratory searchlight RSA analysis. Here, we assessed to which degree neuronal activity is related to either US expectancy or fear ratings, respectively. We then tested whether any brain region is more strongly associated with US expectancy than with fear ratings, but we found no significant results (p < 0.05, FWE corrected). The opposite contrast, fear rating > US expectancy rating, showed widespread results however, with frontal, parietal, and temporal cortex, as well as insula, being more strongly related to fear than to US expectancy ratings (p < 0.05, FWE corrected, Figure 9). This shows that most cortical brain regions we used in the ROI analyses are more closely related to fear ratings than they are related to US expectancy ratings, despite the fact that both types of ratings show largely similar behavioral results.

**Figure 9:**
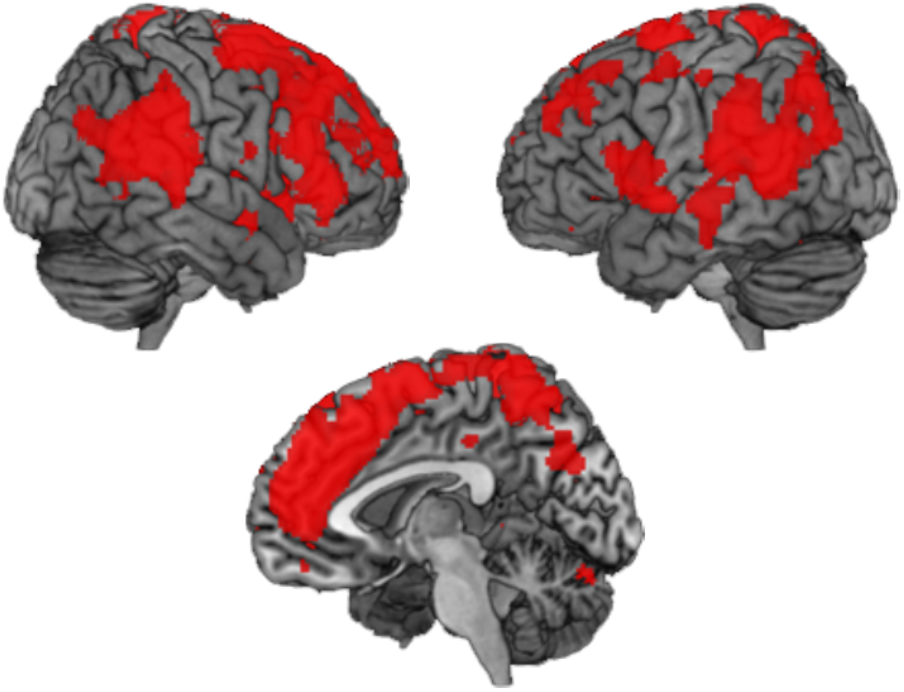
Model-based searchlight RSA results. Brain regions more strongly associated with fear ratings than with expectancy ratings (p < 0.05, FWE corrected)

## 4. Discussion

In this study, our main goal was to investigate the effects of prior CS-US experience on the effects of verbal reversal instructions during fear reversal. We expected verbal reversal instructions to fully explain neural responses after reversal in all brain regions, except the right amygdala. Here, we expected to find evidence for a Pavlovian trace, i.e. a lingering representation of prior experienced CS-US pairings which goes counter to verbal instructions. Using representational similarity analysis, we found that verbal reversal instructions had a profound effect on CS representations. Surprisingly, even though we found strong evidence for an effect of verbal reversal instructions and no evidence for lingering effects of experienced CS-US pairings on behavior, almost all fear-related brain regions included here showed evidence for a Pavlovian trace.

The interaction of associative learning processes and verbal instructions received increased attention in recent years (for a review see Mertens et al., 2018). Much of the past research focused on understanding the effect of verbal instructions on associative learning, showing faster conditioning (Ugland et al., 2013; Atlas et al., 2016) and delayed extinction (Mertens and De Houwer, 2017) for verbally instructed CS-US pairings, highlighting the role of the dACC and dmPFC in verbally mediated fear learning (Mechias et al., 2010). Verbal reversal instructions have been shown to fully reverse acquired fear responses, including startle reflexes (Mertens and De Houwer, 2016). Some initial fMRI evidence further suggests that most fear-related brain regions are susceptible to verbal fear instructions, with the exception of the (right) amygdala (Braem et al., 2017; Atlas et al., 2016). Overall, verbal instructions are a powerful tool to alter previously experienced CS-US associations quickly and efficiently.

Here, we asked the opposite question however: How does prior CS-US experience affect the efficacy of verbal reversal instructions? Previous evidence on psychophysiological measures suggests that reversal effects can be fully explained through verbal instructions only, with no effect uniquely attributable to prior CS-US experience (Mertens and De Houwer, 2016). However, whether similar effects can be replicated for neural coding of fear-relevant stimuli remained unknown.

### 4.1 CS-US experience effects in the right amygdala and beyond

During initial fear learning, we found that both merely instructed and instructed + experienced threats led to robust threat responses, and that each fear-relevant ROI dissociated between these two threatening CSs. After establishing that CS-US experience did affect initial fear learning, we used verbal instructions to reverse CS-US pairings, and tested whether prior experience modulated the efficacy of these instructions. As stated above, for most brain regions we expected verbal instructions to fully reverse CS-US associations, with no residual effect attributable to prior CS-US experience. We hypothesized that the only exception would be the right amygdala, which has a key role in experience-based fear learning more generally (Öhman and Mineka, 2001; Critchley et al., 2002; Knight et al., 2009; Tabbert et al., 2011), shows lingering threat representations for CS+Es in a static environment (Braem et al., 2017), and when CS+Es (but not CS+Is) were reversed multiple times in an experiment (Atlas et al., 2016). Here, we combined the strength of the latter two studies by directly comparing dynamic reversals of CS+E and CS+I within participants (similar to Raes et al., 2014; Mertens and De Houwer, 2016), allowing us to clearly identify any lingering threat representations after verbal reversal instructions were received and to attribute such effects specifically to prior CS-US experience.

Unexpectedly, and counter to previous findings, we found evidence for a Pavlovian trace in almost all brain regions assessed in this study. We found that verbally instructing a CS-US reversal for experienced CSs (CS+E reversed = previously CS+E, but now safe) led to neural representations being more similar to previously experienced and still threatening CSs (= CS+E non-reversed) than to previously instructed and still threatening CSs (= CS+I non-reversed). The latter condition served as a conservative baseline in this comparison, since the only difference between the compared conditions was the experience of CS-US pairings, keeping threat constant. This pattern of results was found in all ROIs assessed here, with the exception of the vmPFC, and more interestingly the right amygdala. Evidence in the latter region was inconclusive though, so that the current data does not allow us to draw strong conclusions about the presence of Pavlovian traces in right amygdala.

One reason for this could be that a small subcortical region like the amygdala showed overall weak signal-to-noise ratios, making detection of reliable voxel pattern responses difficult. Another reason could be our deliberate use of visually distinct but initially functionally identical CSs (e.g., CS+E reversed and CS+E non-reversed). A region that closely monitors, traces, and represents experienced CS-US pairings might benefit more from separable, rather than similar neural pattern responses to these functionally identical CSs. That is, if the right amygdala kept track of different relevant CSs, it could be that it relies more on dynamically changing, separable neural representations, rather than arguably more inflexible, shared representations. Similarly, it has been shown that prefrontal cortex represents two separate items in working memory by using separable, orthogonal representational subspaces, which allows it to efficiently update or select either of them upon demand (Panichello and Buschman, 2021). While it is important to emphasize that this reasoning is post-hoc, this could explain why the right amygdala failed to show a more similar pattern responses for r(CS+E reversed, CS+E non-reversed) than for r(CS+E reversed, CS+I non-reversed). This observation would also explain the different findings in previous studies (Atlas et al., 2016; Braem et al., 2017), which relied on measuring the effect of CS-US experience on neural responses within a single, visually and functionally identical CS.

### 4.2 CS-US experience and instruction validity

While the reasoning above might explain the absence of effects in the right amygdala, the widespread presence of prior CS-US experience effects in other cortical and sub-cortical brain regions is more difficult to reconcile with past findings. Our post-hoc explanation for this unexpected finding is that the current task context might have encouraged participants to rely more heavily on past experiences in evaluating different CS than was the case in previous studies. The design of the merely instructed CS+s (CS+I), which serve as a control condition in this and previous experiments, is critical here. Previously, CS+Is were implemented by replacing aversive USs with a visual placeholder stimulus (Raes et al., 2014; Mertens and De Houwer, 2016; Braem et al., 2017), with participants being instructed that the placeholder will be replaced with the actual US at some point in the experiment. Although this approach controls for the presence of reinforcement on CS+I trials, placeholders have disadvantages as well. They require a cover story, which participants might not believe, and likely induce additional, potentially interfering cognitive processes, such as expectations about how and when they will be replaced by the US. Therefore, we chose to omit placeholder stimuli in our study, and fully rely on verbal threat instructions only. This allowed us to omit cover stories and significantly reduce the complexity of already complex reversal instructions to participants. Additionally, it has been shown that verbal instructions can lead to consistent fear responses even in the absence of placeholders (Ugland et al., 2013), as also suggested by our fear and US expectancy ratings.

However, we believe that this seemingly subtle design choice potentially had significant effects on the weighting of CS-US experience. The instructions were not 100% valid in the beginning of the experiment (because not all CS+s were actually followed by a shock), as well after the reversal instructions (because no more shocks were given in the second reversal phase). If instructions were perceived as an unreliable source about the threat or safety of CSs, this could have led participants to rely more heavily on their prior experiences. Such an explanation would also indicate that what we originally designed as an analysis of “Pavlovian traces”, i.e. lingering effects of prior CS-US experience, instead could be interpreted as an “instruction validity” effect which has little to do with low-level fear-learning mechanisms (Öhman and Mineka, 2001). This could also explain why these widespread effects were also present in cortical brain regions commonly associated with verbal instruction implementation (Demanet et al., 2016; Bourguignon et al., 2018). However, systematic investigations will be needed to directly compare this assumed impact of placeholders on instruction validity and weighting or prior experience.

### 4.3 Conclusion

In sum, we demonstrated that CS-US experience showed no additional effect on self-reported fear and US expectancy ratings after reversal instruction, but substantially affected neural pattern responses, far beyond the amygdala. In relation to previous studies, our findings suggest that the effects of CS-US experience might be co-dependent on the experienced validity of prior instructions, opening up new avenues for future research on experience-instruction interactions in fear reversal learning. Finally, our results have implications for models of fear learning in the right amygdala, bringing important nuance to its role in CS-US experience tracking.

## Acknowledgements

D.W. was supported by was supported by the European Union’s Horizon 2020 research and innovation programme under the Marie Skłodowska-Curie grant agreement no. 665501, the Flemish Science Foundation (FWO, FWO.KAN.2019.0023.01), and the Special Research Fund of Ghent University. S.B. was supported by an ERC Starting grant (European Union’s Horizon 2020 research and innovation programme, Grant agreement 852570). C.G.G. was funded by European Union’s Horizon 2020 research and innovation programme under the Marie Sklodowska-Curie grant agreement no. 835767, and the Spanish Ministry of Science and Innovation, grant ID IJC2019-040208-I. J.D.H. was supported by Ghent University Grant BOF16/MET_V/002. M.B. was supported by an Einstein Strategic Professorship (Einstein Foundation Berlin), and the FWO (project G.0231.13).

## References

Andraszewicz S, Scheibehenne B, Rieskamp J, Grasman R, Verhagen J, Wagenmakers E-J (2015) An Introduction to Bayesian Hypothesis Testing for Management Research. J Manag 41:521–543.

Atlas LY (2019) How instructions shape aversive learning: higher order knowledge, reversal learning, and the role of the amygdala. Curr Opin Behav Sci 26:121–129.

Atlas LY, Doll BB, Li J, Daw ND, Phelps EA (2016) Instructed knowledge shapes feedback-driven aversive learning in striatum and orbitofrontal cortex, but not the amygdala. eLife 5:e15192.

Atlas LY, Phelps EA (2018) Prepared stimuli enhance aversive learning without weakening the impact of verbal instructions. Learn Mem 25:100–104.

Bourguignon NJ, Braem S, Hartstra E, De Houwer J, Brass M (2018) Encoding of Novel Verbal Instructions for Prospective Action in the Lateral Prefrontal Cortex: Evidence from Univariate and Multivariate Functional Magnetic Resonance Imaging Analysis. J Cogn Neurosci 30:1170–1184.

Braem S, Houwer JD, Demanet J, Yuen KSL, Kalisch R, Brass M (2017) Pattern analyses reveal separate experience-based fear memories in the human right amygdala. J Neurosci:0908–0917.

Critchley HD, Mathias CJ, Dolan RJ (2002) Fear Conditioning in Humans: The Influence of Awareness and Autonomic Arousal on Functional Neuroanatomy. Neuron 33:653–663.

Demanet J, Liefooghe B, Hartstra E, Wenke D, De Houwer J, Brass M (2016) There is more into ‘doing’ than ‘knowing’: The function of the right inferior frontal sulcus is specific for implementing versus memorising verbal instructions. NeuroImage 141:350–356.

Etzel JA, Zacks JM, Braver TS (2013) Searchlight analysis: promise, pitfalls, and potential. Neuroimage 78:261–269.

Fullana MA, Harrison BJ, Soriano-Mas C, Vervliet B, Cardoner N, Àvila-Parcet A, Radua J (2015) Neural signatures of human fear conditioning: an updated and extended meta-analysis of fMRI studies. Mol Psychiatry Available at: http://www.nature.com/mp/journal/vaop/ncurrent/full/mp201588a.html [Accessed November 9, 2015].

Knight DC, Waters NS, Bandettini PA (2009) Neural substrates of explicit and implicit fear memory. NeuroImage 45:208–214.

Koban L, Jepma M, Geuter S, Wager TD (2017) What’s in a word? How instructions, suggestions, and social information change pain and emotion. Neurosci Biobehav Rev 81:29–42.

Kriegeskorte N, Mur M, Bandettini P (2008) Representational Similarity Analysis – Connecting the Branches of Systems Neuroscience. Front Syst Neurosci 2 Available at: http://www.ncbi.nlm.nih.gov/pmc/articles/PMC2605405/ [Accessed September 25, 2015].

Lonsdorf TB et al. (2017) Don’t fear ‘fear conditioning’: Methodological considerations for the design and analysis of studies on human fear acquisition, extinction, and return of fear. Neurosci Biobehav Rev 77:247–285.

Lykken DT, Venables PH (1971) Direct Measurement of Skin Conductance: A Proposal for Standardization. Psychophysiology 8:656–672.

Maldjian JA, Laurienti PJ, Kraft RA, Burdette JH (2003) An automated method for neuroanatomic and cytoarchitectonic atlas-based interrogation of fMRI data sets. NeuroImage 19:1233–1239.

Maren S (2001) Neurobiology of Pavlovian Fear Conditioning. Annu Rev Neurosci 24:897–931.

Mechias M-L, Etkin A, Kalisch R (2010) A meta-analysis of instructed fear studies: Implications for conscious appraisal of threat. NeuroImage 49:1760–1768.

Mertens G, Boddez Y, Sevenster D, Engelhard IM, De Houwer J (2018) A review on the effects of verbal instructions in human fear conditioning: Empirical findings, theoretical considerations, and future directions. Biol Psychol 137:49–64.

Mertens G, De Houwer J (2016) Potentiation of the startle reflex is in line with contingency reversal instructions rather than the conditioning history. Biol Psychol 113:91–99.

Mertens G, De Houwer J (2017) Can Threat Information Bias Fear Learning? Some Tentative Results and Methodological Considerations. J Exp Psychopathol 8:390–412.

Nili H, Wingfield C, Walther A, Su L, Marslen-Wilson W, Kriegeskorte N (2014) A Toolbox for Representational Similarity Analysis. PLOS Comput Biol 10:e1003553.

Öhman A, Mineka S (2001) Fears, phobias, and preparedness: Toward an evolved module of fear and fear learning. Psychol Rev 108:483–522.

Olsson A, Phelps EA (2007) Social learning of fear. Nat Neurosci 10:1095–1102.

Panichello MF, Buschman TJ (2021) Shared mechanisms underlie the control of working memory and attention. Nature 592:601–605.

Raes AK, De Houwer J, De Schryver M, Brass M, Kalisch R (2014) Do CS-US Pairings Actually Matter? A Within-Subject Comparison of Instructed Fear Conditioning with and without Actual CS-US Pairings. PLoS ONE 9:e84888.

Riberto M, Paz R, Pobric G, Talmi D (2022) The neural representations of emotional experiences are more similar than those of neutral experiences. J Neurosci Available at: https://www.jneurosci.org/content/early/2022/02/11/JNEUROSCI.1490-21.2022 [Accessed February 25, 2022].

Spielberger CD, Sydeman SJ, Owen AE, Marsh BJ (1999) Measuring anxiety and anger with the State-Trait Anxiety Inventory (STAI) and the State-Trait Anger Expression Inventory (STAXI). In: The use of psychological testing for treatment planning and outcomes assessment, 2nd ed, pp 993–1021. Mahwah, NJ, US: Lawrence Erlbaum Associates Publishers.

Tabbert K, Merz CJ, Klucken T, Schweckendiek J, Vaitl D, Wolf OT, Stark R (2011) Influence of contingency awareness on neural, electrodermal and evaluative responses during fear conditioning. Soc Cogn Affect Neurosci 6:495–506.

Ugland CCO, Dyson BJ, Field AP (2013) An ERP study of the interaction between verbal information and conditioning pathways to fear. Biol Psychol 92:69–81.

Visser RM, Scholte HS, Beemsterboer T, Kindt M (2013) Neural pattern similarity predicts long-term fear memory. Nat Neurosci 16:388–390.

Visser RM, Scholte HS, Kindt M (2011) Associative Learning Increases Trial-by-Trial Similarity of BOLD-MRI Patterns. J Neurosci 31:12021–12028.

Walther A, Nili H, Ejaz N, Alink A, Kriegeskorte N, Diedrichsen J (2016) Reliability of dissimilarity measures for multi-voxel pattern analysis. NeuroImage 137:188–200.

